# Optical activation of TrkB neurotrophin receptor in mouse ventral hippocampus promotes plasticity and facilitates fear extinction

**DOI:** 10.1101/2021.02.14.431126

**Authors:** Juzoh Umemori, Giuliano Didio, Frederike Winkel, Maria Llach Pou, Juliana Harkki, Giacomo Lo Russo, Maarten Verie, Hanna Antila, Chloe Buj, Tomi Taira, Sari E. Lauri, Ramon Guirado, Eero Castrén

**Author notes:** shared corresponding author Juzoh Umemori, Neuroscience center, HiLife, University of Helsinki, 00790 Helsinki, Finland, Eero Castrén, Neuroscience center, HiLife, University of Helsinki, P.O.Box 63 (Biomedicum 1), 00014 Helsinki, Finland. shared first author (Juzoh Umemori, Giuliano Didio).

## Abstract

Successful extinction of traumatic memories depends on neuronal plasticity in the fear extinction network. However, the mechanisms involved in the extinction process remain poorly understood. Here, we investigated the fear extinction network by using a new optogenetic technique that allows temporal and spatial control of neuronal plasticity *in vivo*. We optimized an optically inducible TrkB (CKII-optoTrkB), the receptor of the brain-derived neurotrophic factor, which can be activated upon blue light exposure to increase plasticity specifically in pyramidal neurons. The activation of CKII-optoTrkB facilitated the induction of LTP in Schaffer collateral-CA1 synapses after brief theta-burst stimulation and increased the expression of FosB in the pyramidal neurons of the ventral hippocampus, indicating enhanced plasticity in that brain area. We showed that optical stimulation of the CA1 region of the ventral hippocampus during fear extinction training led to an attenuated conditioned fear memory. This was a specific effect only observed when combining extinction training with CKII-optoTrkB activation, and not when using either intervention alone. Thus, TrkB activation in ventral CA1 pyramidal neurons promotes a state of neuronal plasticity that allows extinction training to guide neuronal network remodeling to overcome fear memories. Our methodology is a powerful tool to induce neuronal network remodeling in the adult brain, and can attenuate neuropsychiatric symptoms caused by malfunctioning networks.

## Introduction

Under pathological conditions, such as post-traumatic stress disorder (PTSD), phobias, and depression/anxiety disorders, traumatic memories are repeatedly and improperly retrieved (1,2). Exposure therapy, where the subject is repeatedly exposed to fear-inducing stimuli under safe conditions, is a widely used method to extinguish or suppress fear responses (3). Fear extinction has been successfully modeled in both humans and animals using the Pavlovian fear conditioning/extinction paradigm, where a neutral conditioned stimulus (CS, tone or context) starts to elicit a fear response after being associated with an aversive unconditioned stimulus (US). This fear response is reduced after a repeated exposure to the CS without the US (4–6). Although extinction training gradually reduces the fear responses in human patients and adult rodents, these fear responses tend to reappear with time or upon later re-exposure to the CS, a phenomena known as “spontaneous recovery”, and “fear renewal” when induced by a neutral cue or the same context, respectively (7,8). We have previously shown that the combination of extinction training and chronic treatment with fluoxetine, a commonly used antidepressant, but neither treatment alone, induces an enduring loss of conditioned fear memory in adult mice (9), which is similar to the permanent fear extinction found in early postnatal mice (10,11). A chronic treatment with fluoxetine reactivates a state of plasticity similar to that observed during the critical periods of plasticity or induced juvenile-like plasticity, a state we refer as iPlasticity, which have been shown in different brain regions, such as the amygdala, medial prefrontal cortex (mPFC), and hippocampus (9,12,13). These observations suggest that fear extinction is a process dependent on the reshaping of neural networks through experience-dependent plasticity. However, the mechanisms through which neural networks are reconfigured are still unknown.

Brain-derived neurotrophic factor (BDNF), through activation of its neurotrophic receptor tyrosine kinase B (TrkB), is thought to be a key factor in neuronal plasticity and required for the iPlasticity by fluoxetine treatment (9,14). The binding of BDNF to TrkB causes dimerization and autophosphorylation of TrkB, leading to activation of intracellular signaling pathways involved in neuronal differentiation, survival, and growth as well as synaptic plasticity in neurons (15,16). These pathways also regulate gene transcription and long-term potentiation (LTP) (17,18). Interestingly, Chang et al created a photoactivatable TrkB (optoTrkB), where full-length TrkB is conjugated with a photolyase homology region (PHR) that dimerizes in response to blue light (470 nm) (19). They have shown that light stimulation can activate the canonical Trk signaling pathways through optoTrkB in a reversible manner, and a prolonged patterned stimulation induces differentiation of cultured neurons (19).

Here, we studied whether activation of TrkB through optoTrkB *in vivo* is sufficient to induce plasticity in the fear circuit and to facilitate fear extinction. For an efficient expression of optoTrkB in pyramidal neurons, we constructed a lentivirus that expresses optoTrkB (19) modified in the following points: (i) optimization of codons of the PHR domain for higher expression in rodents, (ii) attachment of a flexible tag (20) between TrkB and PHR, which allows a better interaction between optoTrkB C-terminus and its partners (21), and (iii) expression of a fusion protein by a short-type (0.4 kb) promoter of calcium/calmodulin-dependent protein kinase type II alpha subunit (CKII) for specific expression in pyramidal neurons (22). After confirming that CKII-optoTrkB lentivirus is expressed and activated in cultured cortical neurons, we activated CKII-optoTrkB in the projection neurons of the ventral hippocampus (vHP), which are known to be involved in fear extinction (23,24) through the modification of mood and spatial memory (25), and conducted the Pavlovian fear conditioning paradigm.

## Material and Method

All animal experiments followed the Council of Europe guidelines and were approved by the State Provincial Office of Southern and Eastern Finland. Detailed procedures are described in the Supplemental Information.

### Mice

C57BL/6JHss were originally purchased from Harlan (Netherlands); 10-to 12-week-old mice were used for this study. Mice were kept under standard laboratory conditions (21°C, 12-h light-dark cycle, light at 6AM) with free access to food and water.

### Infection of lentivirus and optic stimulation of optoTrkB in cultured cortical neurons

Rat primary cortical cultured neurons from E17 rat embryos were prepared using a method reported previously (26). The cells were infected with CKII-optoTrkB lentivirus (initial stock titer 8.37×10^7^ pg/ml [p24]) at day *in vitro* 3 (DIV3) for immunoblotting, while the other cells on coverslips were infected at DIV9 for morphological analyses. The plates were kept in darkness at all time after the infections. The cells were exposed to blue light (LED devices, Mightex) at DIV10 and DIV17 for immunoblotting and immunocytology, respectively. The cells were photo-stimulated 12 times for 5 seconds with a 1-minute inter-trial interval, aiming to mimic the behavioral experiments. The cells were collected immediately for immunoblotting and 24 hours later for immunocytology. For immunoblotting, cells were lysed following a protocol described previously (27) and stored in darkness at −80°C. The samples for immunocytology were fixed with 4% of paraformaldehyde (PFA) and stored in PBS containing 0.02% NaN_3_ at 4 °C.

### Immunoblotting of lysate from cultured cells

Immunoblotting was conducted according to a method reported previously (27). Details on primary and secondary antibodies are provided in supplemental table 1.

Chemiluminescent signals were developed by ECL plus (ThermoFisher Scientific) with a 5-minute incubation according to instructions provided by the manufacturer and detected by a LAS-3000 dark box (Fujifilm).

### Imaging analyses on dendrites after immunocytology

We compared the number of spines in the secondary dendritic branches as described previously (28). All antibodies used for immunocytology are listed in supplemental table 1. The stained cells were imaged with a Leica TCS SP8 X with a magnification of 40x for analysis of primary neurites and spines. The primary branches sprouting from the soma were counted blindly and manually. Spines on the second branches were randomly and blindly selected. The number and type of spines were analyzed manually.

### Lentivirus infection and implantation of optic cannulas

Mice were anesthetized with Isoflurane and fixed on a stereotactic frame. A total of 1 μl of virus solution was injected into the vHP, at 3.1 mm caudally and +/- 2.0 mm laterally from the Bregma with a depth of 3.9 mm and an angle of 18°. Three weeks after infection, optic fibers with cannula were inserted into the vHP, 3.1 mm caudally and +/- 3.0 mm laterally from the Bregma, with a depth of 3.0 mm and an angle of 4° and fixed and sealed with dental cement (Tetric Evo Flow). The mice were kept in single cages and underwent the fear conditioning/extinction test 1 week after the implantation.

### Electrophysiology

Field excitatory postsynaptic potentials (fEPSP) were recorded in acute hippocampal slices. Briefly, the brains were isolated 4 weeks after infection with CKII-optoTrkB and immediately immersed in ice-cold dissection solution (29), after which 350-μm brain slices were cut and incubated for recovery for 45 minutes at 31 to 32°C in artificial cerebrospinal fluid (ACSF) (30). The slices were stimulated by light (LED 480 nm) three times for 5 seconds every minute. fEPSPs were then recorded in an interface chamber using ACSF-filled glass microelectrodes (2-4 MΩ) positioned within the CA1 stratum radiatum in response to Schaffer collateral stimulation (0.05 Hz). Stimulation intensity was adjusted such that the baseline fEPSP slope was 20-40% of the maximal intensity that resulted in the appearance of a population spike. LTP was subsequently induced by tetanus stimulation (100 pulses at 50 Hz) or brief theta-burst stimulation (1 episode of TBS consisting of 2 stimulus trains at 5 Hz with 4 pulses at 100 Hz).

### Fear extinction test with optic stimulation

The fear conditioning paradigm was conducted following a protocol described previously (9). Briefly, the mice were put into Context A and received an electric foot shock (0.6 mA) after a 30-second sound cue (“beep” sounds 80 dB), which was repeated four times with a 30-to 60-second interval. Two days later the mice were put in Context B and received only the sound cue (30s “beep” sound 80 dB) immediately followed by optical stimulation for 5 seconds. The light was applied manually by a single-color LED (470nm wave length) device (Mightex) connected to a BioLED light source Control Module (Mightex), which in turn was connected to optic wires splitting into two parallel optic fibers (Kyocera Inc.) ending on both sides on the ferrules implanted in the head of the mouse. Mice were made able to move freely through a rotary joint (Mightex). The extinction training with light stimulation was repeated 12 times with different intervals (25-60 seconds) for 2 days. One week after, the mice were tested in Context B (spontaneous recovery) followed by exposure to Context A (fear renewal) with the same sound cue (presented 4 times/test) provided during conditioning and extinction. Spontaneous recovery and fear renewal were tested again 3 weeks later as an estimate of remote memory. The durations of freezing were measured as an index of conditioned fear.

### Immunohistochemistry and image analysis on FosB intensity

Mice were infected with CKII-optoTrkB and we implanted optic cannulas into the vHP as described above. The vHP were exposed to light 12 times for 5 seconds after a 30-second sound with different intervals (25-60 seconds) in the same way as in the extinction training through optic cannulas for 2 days. Twenty-four hours after the last stimulation, the animals were perfused transcardially with PBS followed by 4% PFA in PBS. Isolated brains were post-fixed overnight and stored in PBS with 0.02% NaN_3_ until cut on a vibratome (VT 1000E, Leica). Free-floating sections (40 μm) were processed for fluorescence immunohistochemistry following a protocol described previously (31) using the antibodies listed in supplemental table 1. Images were obtained with a Zeiss LSM 710 confocal microscope with a magnification of 20x. The intensity of delta FosB was analyzed by Fiji software (32).

### Statistics

Biochemical data were analyzed by unpaired t-test following F-test. If standard deviation of the two groups were not equal, Welch’s correction was applied. For comparisons of more than two groups, we used one-way ANOVA followed by Holm-Sidak’s multiple comparisons test. Behavioral data was analyzed by two-way ANOVA, taking sessions and light exposure/non-exposure as independent factors, followed by Fisher’s LSD test. All statistical analyses were performed using Prism 6 or 8 (GraphPad Software), and shown in supplemental table 3. A *p*-value <0.05 was considered statistically significant.

## Results

### Construction of CKII-optoTrkB

We optimized the codons of the PHR domain and the resulting Codon Adaptation Index (CAI) (33) was increased to 0.86 in the optimized codon, compared to 0.79 as in Chang’s original PHR (Supplemental fig. 1). The homology of DNA sequences between our construct and the original optoTrkB was 78% (Supplemental fig. 2). The optimized PHR region, flexible tag (20), and full-length TrkB were sub-cloned into a lentivirus backbone vector with a short-type (0.4 kb) CaMKIIa promoter (pFCK(0.4)GW) (22). The CKII-optoTrkB construct was used for lentivirus production (see supplemental note).

### Optical stimulation of optoTrkB activates TrkB signals and neural plasticity in vitro

We determined the optimal virus concentration by testing different concentrations of CKII-optoTrkB in cultured cortical neurons and performed immunoblotting for phosphorylated TrkB (at tyrosine 706 residue, pY706), TrkB itself, and phosphorylated and non-phosphorylated Extracellular signal-regulated kinase (ERK), a downstream signal of the BDNF/TrkB pathway (Supplemental fig. 3). Immunoblotting with optimal concentration of the lentivirus showed an effect of light exposure on phosphorylation of optoTrkB at pY515, pY706, and pY816 (Fig. 2a, b, c, and d). As expected, phosphorylation of endogenous TrkB was not influenced by light (Supplemental fig. 4), but increased only after BDNF treatment (Supplemental fig. 5). Then we verified the phosphorylation of downstream signals of the BDNF/TrkB pathway and observed increased phosphorylation of cAMP response element-binding protein (CREB) and pERK after light stimulation when compared to the control group transfected with optoTrkB but not exposed to light (Fig. 2e, f).

**Figure 1.**
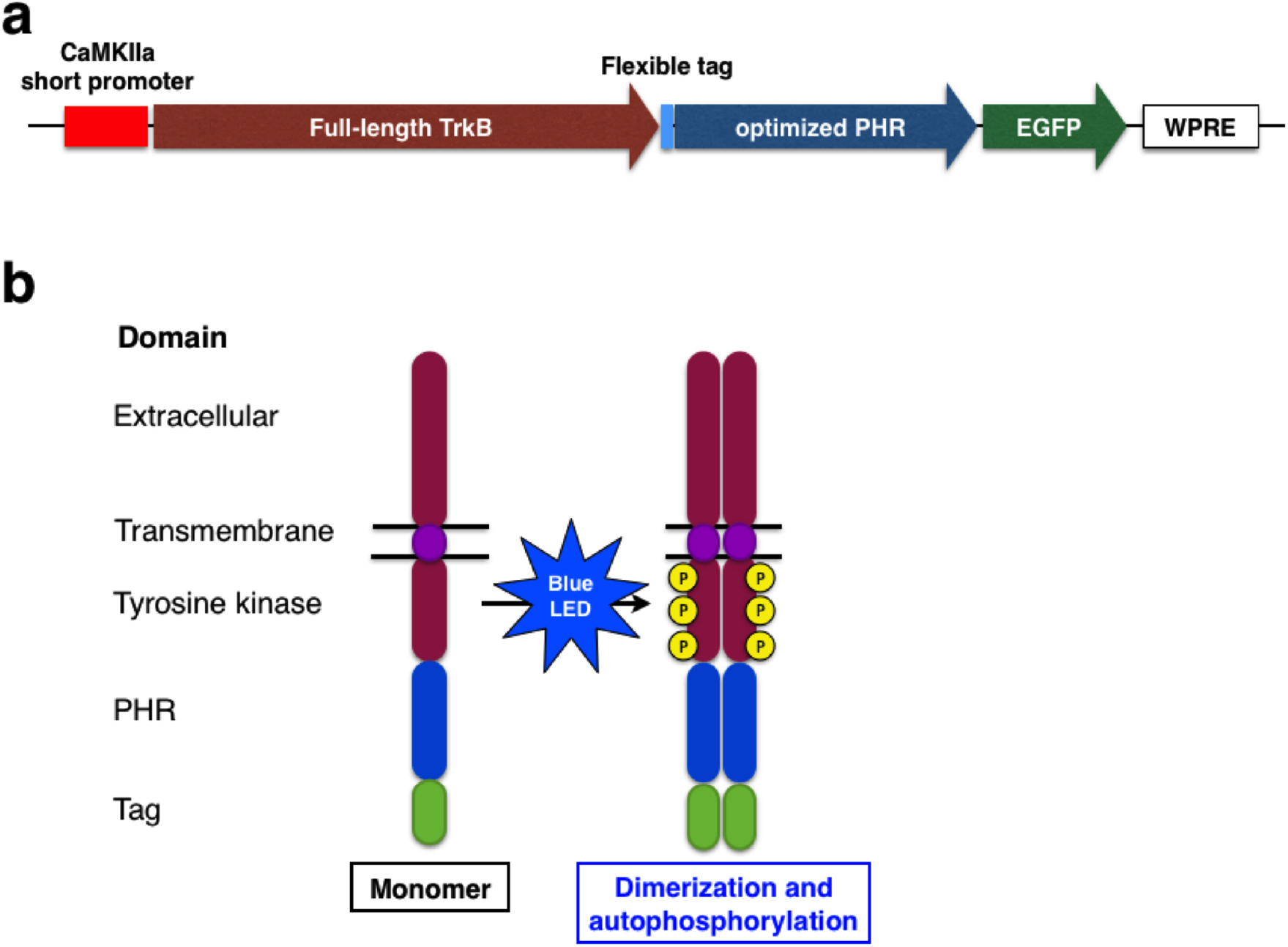
Development of optoTrkB for the *in vivo* study. (a) Gene structure of optoTrkB. CKII-optoTrkB consists of a short version (0.4 kb) of CaMKIIa promoter, full length TrkB, flexible tag, EGFP, and Woodchuck Hepatitis Virus (WHP) Posttranscriptional Regulatory Element (WPRE). (b) Protein structure of optoTrkB. TrkB consists of extracellular, transmembrane, and tyrosine kinase domains and was conjugates with PHR and GFP. The PHR domain dimerizes in response to blue light (470 nm) therefore inducing dimerization and autophosphorylation of TrkB to activate the canonical TrkB signaling pathways.

**Figure 2.**
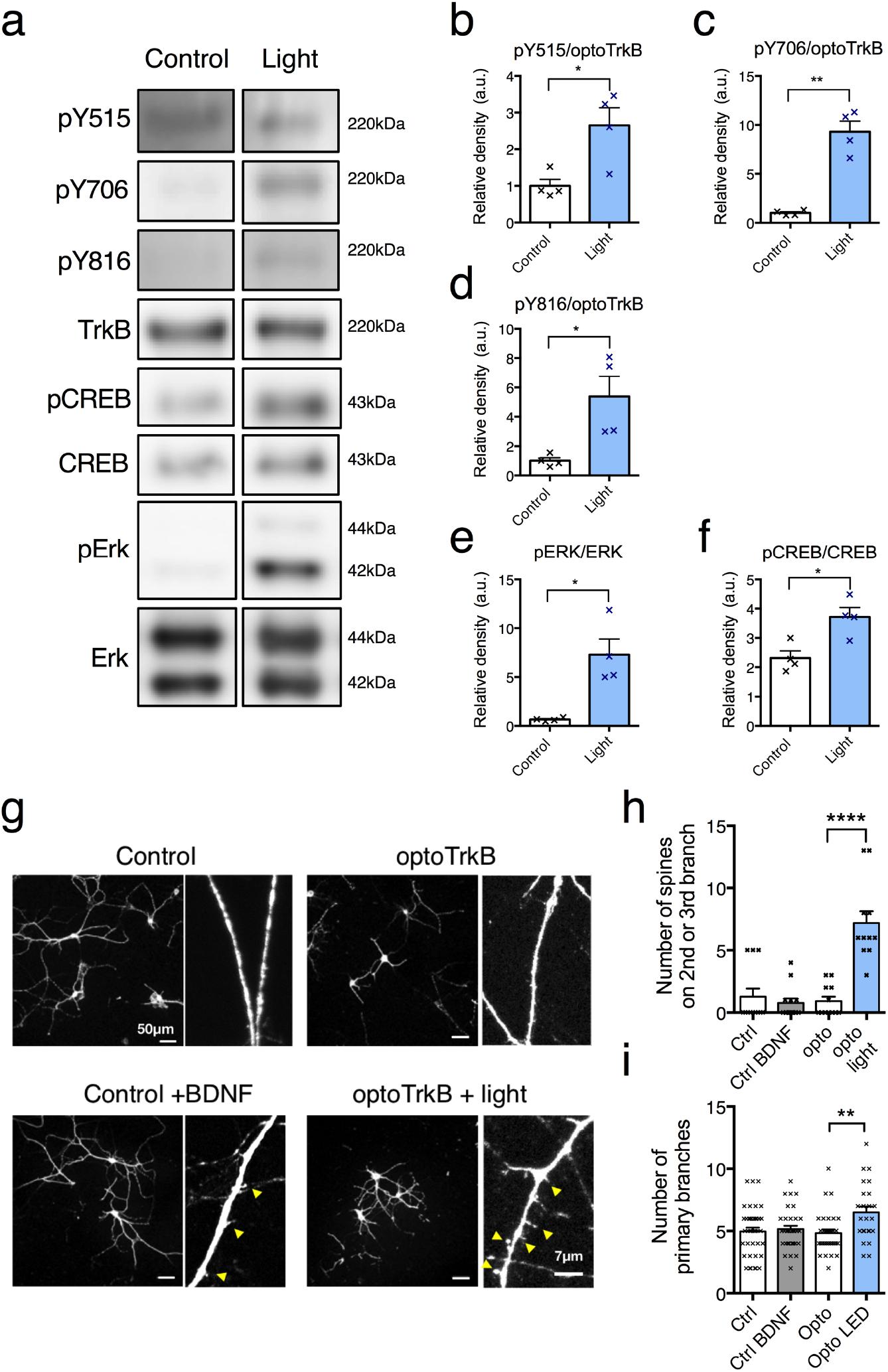
(a) Immunoblotting with antibodies against phosphorylated TrkB and downstream signals of TrkB. Quantitative analysis of phosphorylation of optoTrkB at pY515 (b), pY706 (c), and pY816 (d) phosphorylation site after light stimulation (12 times for 5 seconds with 1-minute interval) (N = 4, each group). The intensity of optoTrkB (220 kDa) was normalized with the non-phosphorylated version of the same protein. There were significant effects of light exposure on phosphorylation of optoTrkB at pY515 (unpaired t-test, p = 0.0178), pY706 (p = 0.004), and pY816 (p = 0.0477). Quantitative analysis of phosphorylation of ERK (e) and CREB (f) after light stimulation. Intensity of the bands of phosphorylated ERK (pERK) and CREB (pCREB) was normalized by non-phosphorylated ERK(42 kDa) and CREB(43 kDa), respectively. The ratios significantly increased after light stimulation (unpaired t-test, pCREB/CREB, p = 0.0133; pERK/ERK, p = 0.025). (g-i) Effects of activation of CKII-optoTrkB on primary dendrites and spines in cultured cortical neurons (DIV17). The uninfected neurons were treated with 5 ng/ml of BDNF as control. The number of spines on 2nd or 3rd branch and primary dendrites extending from the cell body was counted manually. (g) Representative images of MAP2 immunostaining of cortical neurons. (h) The number of spines was not increased after a 5ng/ml BDNF treatment (Holm-Sidak’s multiple comparisons test, Control vs Control + BDNF, p = 0.6898). However, the number of spines after activation of CKII-optoTrkB was significantly higher than non-activated infected cells (one-way ANOVA, p < 0.0001; Holm-Sidak’s multiple comparisons, optoTrkB vs optoTrkB light, p < 0.0001 (N=11-13 in each group). (i) The number of primary dendrites was not increased after BDNF treatment (Holm-Sidak’s multiple comparisons test, Control vs Control + BDNF, p = 0.6898). However, the number of spines after activation of CKII-optoTrkB was significantly higher than non-activated infected cells (one-way ANOVA, p = 0.0053; Holm-Sidak’s multiple comparisons, optoTrkB vs optoTrkB light, p = 0.0023 (N=11-13 in each group).. (N = 48-63 in each group). CREB, cAMP response element-binding protein; ERK (Extracellular signal-regulated kinase). Scale bar, 50 μm. Bars represent means ± SEM. * p < 0.05, ** p < 0.01, *** p < 0.001.

To verify if the activation of optoTrkB causes morphological changes of dendrites and spines *in vitro*, we stained cortical primary neurons with MAP-2 antibody (Fig. 2g). The number of spines on second- and third-order dendritic branches was increased at 24 hours after light stimulation but not after BDNF treatment (5ng/ml) (Fig. 2h). Furthermore, we found an increased number of primary dendrites extending from the cell body in light-stimulated CKII-optoTrkB infected cells compared to non-stimulated cells or BDNF-treated cells. These results indicate that activation of optoTrkB promotes initial neurite and spine formation more rapidly than the BDNF treatment (28).

### Increased FosB expression after activation of optoTrkB in the ventral hippocampus

To confirm the activation of optoTrkB after light stimulation, optoTrkB lentivirus-infected mice were perfused 24 hours after 2 days of optic stimulation (12 sessions of 5-second exposure). Immunohistochemistry showed an increase of delta FosB expression in the regions close to the infection sites in the CA1 of the vHP (Fig. 3a and b), indicating that optoTrkB promotes BDNF/TrkB signals, as shown previously in case of overexpression of BDNF (34).

**Figure 3.**
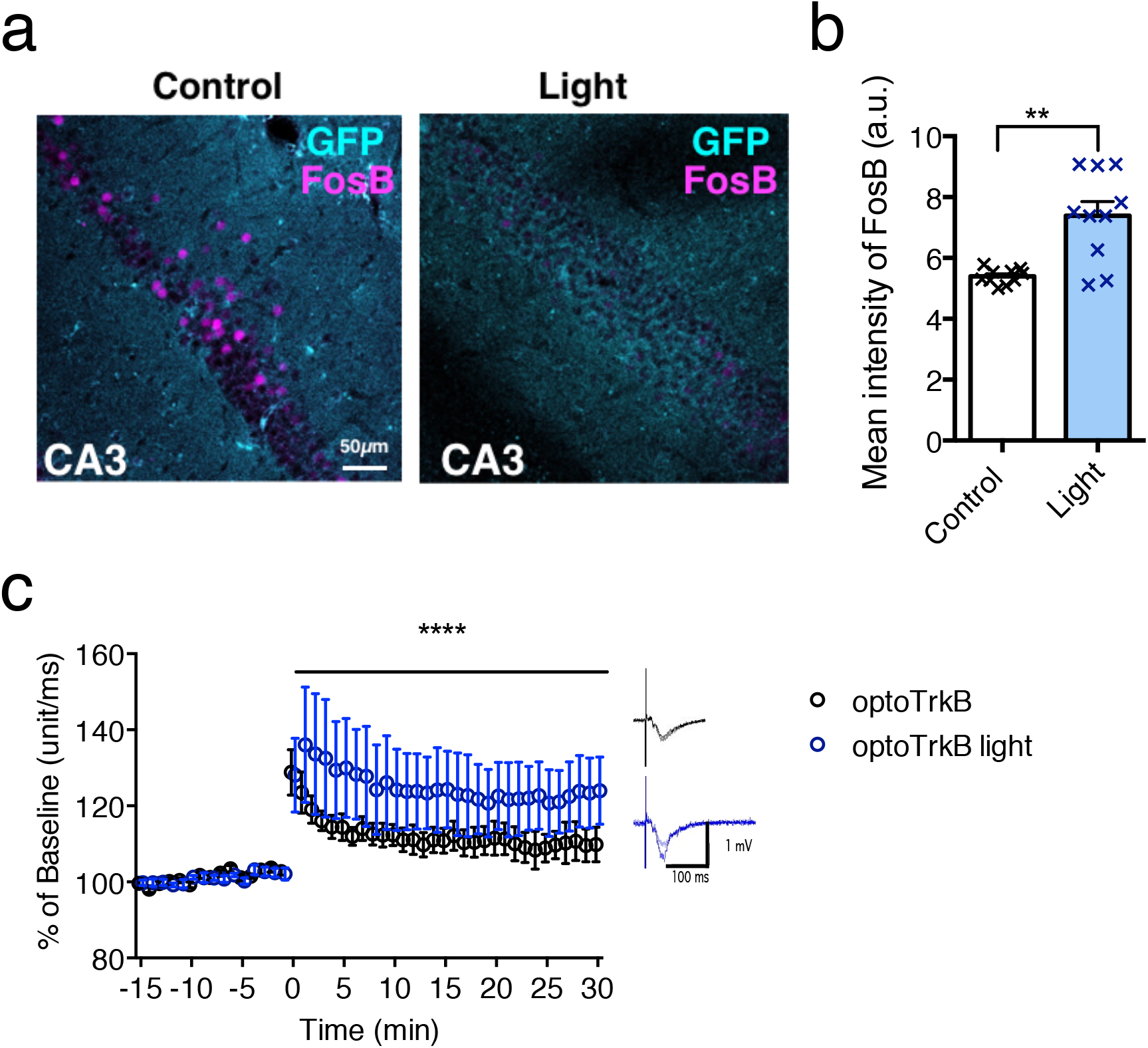
Enhanced neural plasticity after optoTrkB activation ex vivo. BDNF/TrkB signals were activated in the pyramidal neurons in the CA1 of the vHP after optic stimulation of CKII-optoTrkB. (a) Representative figures of the delta FosB and EGFP stainings close to the infection sites in the CA1of vHP. (b) Comparison of the intensity of delta FosB staining between non-light and light stimulation (N = 10, each group). Delta FosB immunoreactivity was higher in the group with light stimulation compared to the group without light stimulation (unpaired t-test, p = 0.0006). (c) The slices from CKII-optoTrkB infected mice were activated by light for 30 seconds. After 30 minutes, long-term potentiation (LTP) was induced by brief theta burst. fEPSPs during the last 10 minutes of recording were significantly larger in the group with light exposure compared to the group without light (two-way ANOVA, p < 0.0001). Pictures in the right panel show representative traces of fEPSC during baseline and after LTP induction. (optoTrkB, N = 5; optoTrkB light, N = 7). Error bars indicate mean ± SEM.

### LTP is potentiated after activation of optoTrkB

To investigate synaptic plasticity in the hippocampal circuitry, fEPSPs were recorded in acute hippocampal slices obtained from CMKII-optoTrkB lentivirus-infected mice after *ex vivo* light stimulation. A traditional tetanus stimulation resulted in a comparable induction of LTP in control and light-stimulated slices (Supplemental Fig. 6), suggesting that a strong tetanization induces LTP independently from optoTrkB activation. However, a brief theta-burst stimulation led to a robust increase in synaptic strength only in light-stimulated slices (Fig. 3c), while it produced a slight potentiation of fEPSPs in slices infected with optoTrkB without light exposure (control). This indicates that the activation of CKII-optoTrkB facilitates LTP induced by a brief theta-burst stimulation in the hippocampus.

### Activation of optoTrkB during fear extinction training reduces fear memory

Since the vHP is thought to be a key brain region for the processing of the extinction of contextual fear memory (23), we hypothesized that the activation of optoTrkB in the vHP during fear extinction may promote fear erasure. To test this hypothesis, we performed the fear conditioning paradigm (Fig. 4a). CKII-optoTrkB lentivirus was infected bilaterally into the CA1 region of the vHP, and optic cannulas were implanted into the same region (see Material and method) (Fig. 4b). During fear-conditioning/acquisition, all infected and implanted mice were conditioned by exposing them to a mild foot shock paired with a sound cue in context A, and all mice showed increased freezing (supplemental fig. 6). The mice were then equally divided into the following two groups: control (without stimulation) and light exposure (supplemental fig. 6). Two days later, the vHP was bilaterally exposed to light through optical fibers for 5 seconds immediately after the CS (“beep” sounds) in context B during 2 days of extinction training. During the extinction training of the first day (Ext1) (Fig. 4c) and the second day (Ext2) (Fig. 4d), both groups showed decreased freezing, but the effect was significantly more pronounced after LED stimulation. One week later there was no difference of freezing in context B (spontaneous recovery) and a weak decrease of the fear renewal in the LED group in context A (fear renewal) (supplemental fig. 7). However, three weeks later, the previously light-stimulated mice showed a decrease of freezing in the spontaneous recovery test (Fig. 4e) and a strong decrease in the fear renewal test (Fig. 4f). These results indicate that the light-stimulated mice initially retain a high freezing representation, but they then reduce the long-term or remote contextual fear memory. CKII-optoTrkB-infected mice stimulated by light without extinction training did not show differences compared to the control group during remote spontaneous recovery (Fig. 4g) or remote fear renewal (Fig. 4h). On the contrary, it was significantly different to the group exposed to both LED and extinction training (Supplemental figure 7), indicating that conditioned fear is reduced only when combining CKII-optoTrkB activation and extinction training but not with either intervention alone.

**Figure 4.**
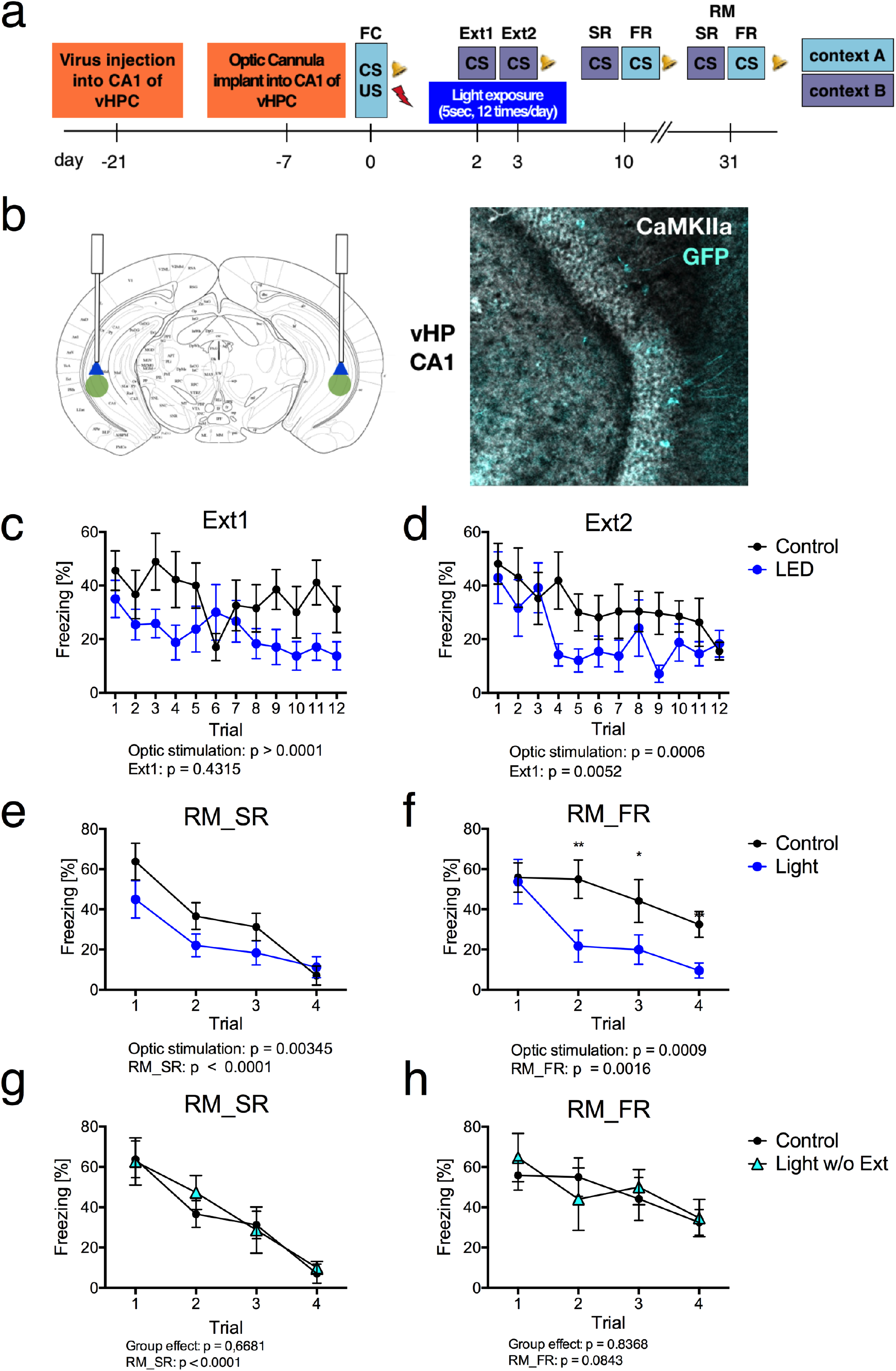
Activation of CKII-optoTrkB combined with extinction training promotes extinction of conditioned fear. (a) Scheme of fear extinction paradigm. Mice were subjected to fear extinction training in context B two days after fear conditioning with tones in context A. After 1 week, mice were then subjected to spontaneous recovery (SR) in context B and fear renewal (FR) in context A. Further 3 weeks later, the mice were subjected again to SR and FR for testing remote memory (RM). (b) CKII-optoTrkB lentivirus was infected into CA1 of the vHP (3.1 mm caudal, ±2 mm latera from Bregma, 3.9 mm depth from *dura mater*) (left). A representative infected site in the vHP (right). (c) (d) Significant effects of sessions were detected in freezing response during day 2 of extinction training (two-way ANOVA, p = 0.0052) and an effect of light stimulation during both days (Day1, p < 0.0001; Day2, p = 0.0006). In addition, there were significant differences between the first extinction session and the session with the lowest freezing duration post hoc (Fisher’s LSD): control, 1st in 1st day vs 12th in 2nd, p = 0.0060; light, 1st in 1st day vs 9th in 2nd, p = 0.0159). Light-stimulated mice showed a decrease of freezing in remote spontaneous recovery (RM_SR) (two-way ANOVA: p = 0.00345) (e) and a strong decrease in remote fear renewal (RM_FR) (p = 0.0009) (f). Post-hoc analysis showed significant differences between two groups after second sessions (Fisher’s LSD: light2-4, p = 0.0002; post-hoc: 2nd, p = 0.0047; 3rd, p= 0.0362; 4th, p = 0.0465). The group with light stimulation but no extinction training (LED w/o Ext) did not show decreased freezing during (g) remote spontaneous recovery (two-way ANOVA, p = 0.6681) or (h) remote fear renewal (two-way ANOVA, p = 0.8368) compared to control. Control (optoTrkB infected without light), N = 8, optoTrkB light, 8; optoTrkB light w/o extinction, 5). Error bars indicate mean ± SEM. * p < 0.05, ** p < 0.01.

## Discussion

We developed a CKII-optoTrkB lentivirus by modifying the original optoTrkB (19) and demonstrated that it can be used to promote plasticity for *in vivo* studies. The optical activation of optoTrkB in the pyramidal neurons in the CA1 of the vHP promotes plasticity in the fear circuitry and enables fear extinction training to greatly reduce the conditioned fear memory, specifically the contextual memory. This study directly demonstrates that the activation of TrkB can promote plasticity-related behaviors in the fear circuitry. Moreover, our optoTrkB approach represents a new system to control plasticity temporarily and spatially.

### Strong and rapid effects of optoTrkB activation compared to BDNF treatment

In cultured cortical neurons, light stimulation of CKII-optoTrkB promoted phosphorylation of CREB and ERK to the same extent as BDNF stimulation, suggesting that activation of CKII-optoTrkB has biochemically comparable effects to BDNF treatment at concentrations of 5 ng/ml. Moreover, in contrast to BDNF treatment, activation of optoTrkB promoted initial neurite and spine formation. It has been reported that a higher BDNF concentration is needed to observe such a drastic increase in the number of primary dendrites and spines (28,35,36). Thus, the activation of CKII-optoTrkB acts as rapidly and efficiently as longer treatment with high concentration of BDNF.

### Neural plasticity is increased after activation of CKII-optoTrkB ex vivo

Previous studies with deleted and mutated TrkB showed impaired LTP at CA1 hippocampal synapses and impaired learning behaviors (15,18), indicating that TrkB activation is critical for LTP induction in the hippocampus. We now demonstrate that after direct activation of TrkB through CKII-optoTrkB a brief TBS produced a robust LTP that was significantly larger compared to controls. We used a modified and “brief” TBS, since a stronger TBS protocol robustly induces LTP in the hippocampus (37). Interestingly, induction of LTP in response to a strong tetanic stimulation was not affected by activation of CKII-optoTrkB, suggesting that TrkB activation lowered the threshold for LTP induction. Alternatively, LTP induced by tetanic stimulation could be reaching a “saturated point”, occluding any facilitation by optoTrkB activation. TBS has been shown to reflect physiological conditions (38). Our results strongly suggest that activation of optoTrkB can sensitize pyramidal neurons in the hippocampal network to be more plastic and to respond to a brief stimulation and adapt to external stimuli, such as an extinction training.

### Activation of optoTrkB combined with extinction training reduces fear memory but does not delete it entirely

In the current study, light-stimulated mice did not completely erase the fear response in the first session, but rather in the second session of the remote fear renewal tests. Similar effects, where fear response is decreased after the second session in fear renewal, were found after PNN removal in the basolateral amygdala (BA) before extinction training (11). In contrast, the combination of extinction training and chronic treatment with fluoxetine decreases the fear response almost completely even in the first session (9). These results suggest that optoTrkB activation in combination with extinction training does not completely replace or erase the conditioned fear memory but it reduces it by promoting neural plasticity in the pyramidal neuron network of vHP during the extinction training. These observations might support the idea that the original fear memory is preserved and the extinction training simply adds a new inhibitory association rather than erasing the original memory (39–41).

### Fear extinction circuitry

Our results suggest that plasticity in pyramidal neurons in the vHP is a key element for processing the extinction of contextual fear memory. Fear extinction is thought to be controlled by a distributed network, including the amygdala, mPFC, and hippocampus (23). Prior evidence suggests that fear memories are disrupted with an increased activity of the vHC; optical activation of these neurons was reported to induce fear extinction and modification of behavior related to mood and spatial memory (25). The vHP may modulate emotional regulation, whereas dorsal HP is thought to contribute to cognitive functions such as learning and memory (42–46). The CA1 region of the vHP in particular sends strong projections to other regions, such as the BA, hypothalamus, nucleus accumbens, and the mPFC (47–50), and there is evidence that these projections process emotional behavior (51–53). In addition, impaired function in Hippocampal-prefrontal circuit has been observed in psychiatric patients including PTSD and schizophrenic subjects (54), and configurational changes in prefrontocortical inputs from Amygdala and Hippocampus have been suggested as a possible mechanism underlying psychiatric disorders (55). Furthermore, it has been reported that BDNF infusion into the PFC and HP erases fear memory (56), and engram cells of projection neurons in CA1 of vHC play a necessary and sufficient role in social memory (57). Recently, Jimenez et al demonstrated that optogenetic activation of the CA1 terminals in BA impaired contextual fear memory (53). Thus, our results suggest that plastic changes in the projection neurons in CA1 of the vHP, enabled by optoTrkB activation, modify anxiogenic contextual information when combined with fear extinction training.

### iPlasticity and application of optoTrkB system in vivo

Many kinds of interventions induce iPlasticity, where networks in adult brain are allowed to better adapt to the changes in the internal and external milieu (16,17,58). In addition to fear extinction, specific training and other external manipulations, when combined with fluoxetine, have been shown to increases neural plasticity and alter symptoms of neuropsychiatric diseases in models such as ocular dominance plasticity (14) and socialization animal models (59). We hypothesize that these effects are modulated via the BDNF/TrkB pathway, but it is not yet clear which neural pathways are modified through iPlasticity in these behaviors. OptoTrkB is a new tool for controlling neural plasticity in a temporal and cell-type specific manner and the optic control of neural plasticity adds another dimension to traditional optogenetics using direct activation and inhibition of neurons by channelrhodopsin and halorhodopsin in experimental neurosciences.

## ACKNOWLEDGMENTS

We thank Sulo Kolehmainen and Outi Nikkila for assistance in all experiments. We also thank caretakers in the F-building in UH for assistance with animal care, and Lakovos Lazaridis and Konstantinos Meletis in the Karolinska institute for kindly providing instruction in optogenetic techniques. The original research in our laboratory was supported by the ERC grant # 322742 – iPLASTICITY, the Sigrid Jusélius foundation, the EU Joint Programme–Neurodegenerative Disease Research (JPND) project # JPCOFUND_FP-829-007, the HiLife Fellows program, the Academy of Finland grants #294710, 303124, and 307416, Bilateral exchange program between the Academy of Finland and JSPS (Japan Society for the Promotion of Science), the Brain & Mind grants, and the University of Helsinki Research Foundation.

## AUTHOR CONTRIBUTIONS

J.U., R.G., and E.C. conceived of and designed the project. J.H., G.D., G.L., MV, and J.U. performed experiments related to cultured cells. M.L., H.A., G.D, F.W., and J.U. conducted operations, behavioral experiments, and immunohistochemistry. F.W. performed electrophysiological experiments under the supervision of T.T. and S.L. All authors were involved in writing the manuscript.

## Supplemental figures

**Supplemental Figure 1.**
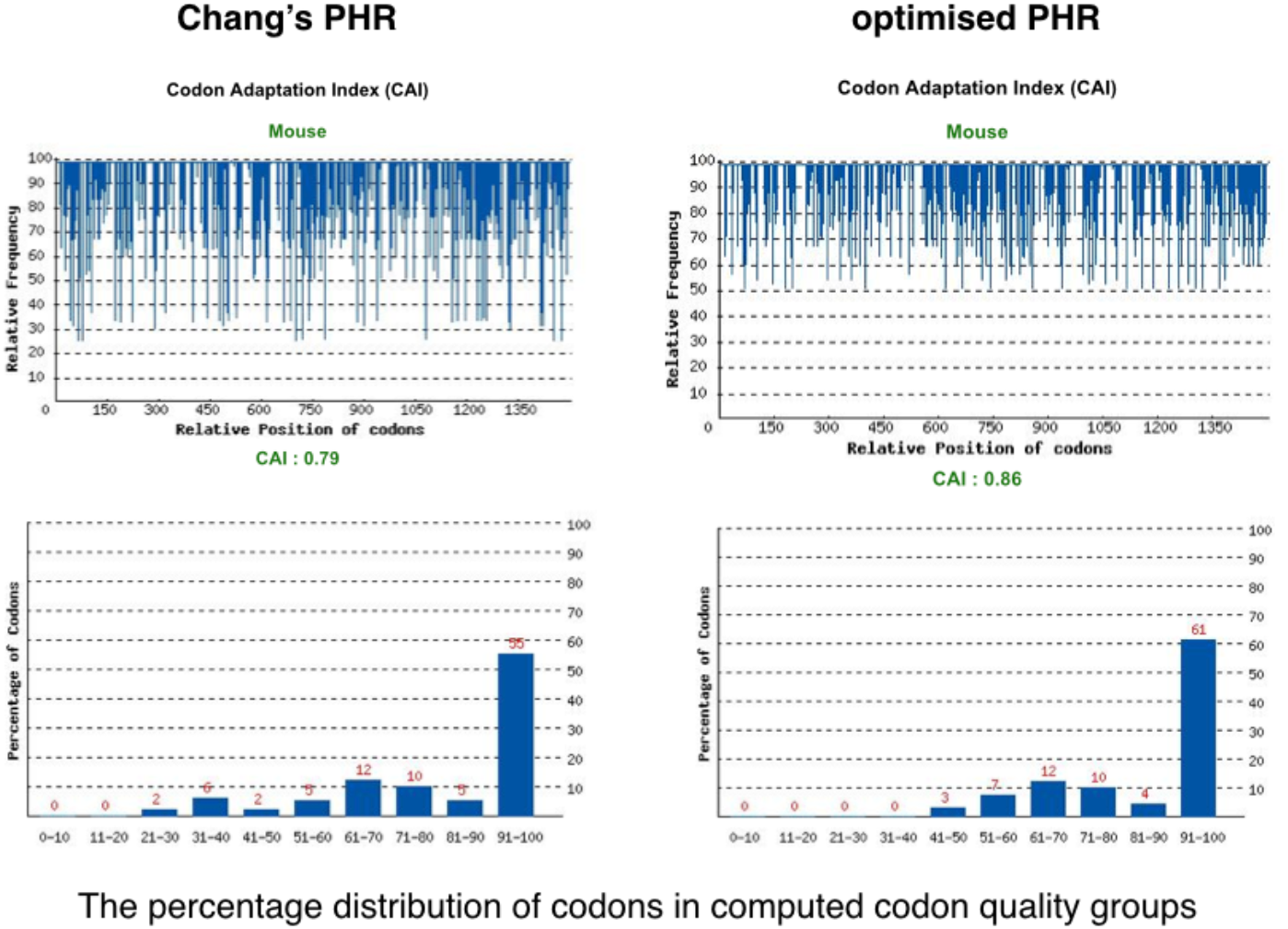
optimized codons of PHR domain

**Supplemental Figure 2.**
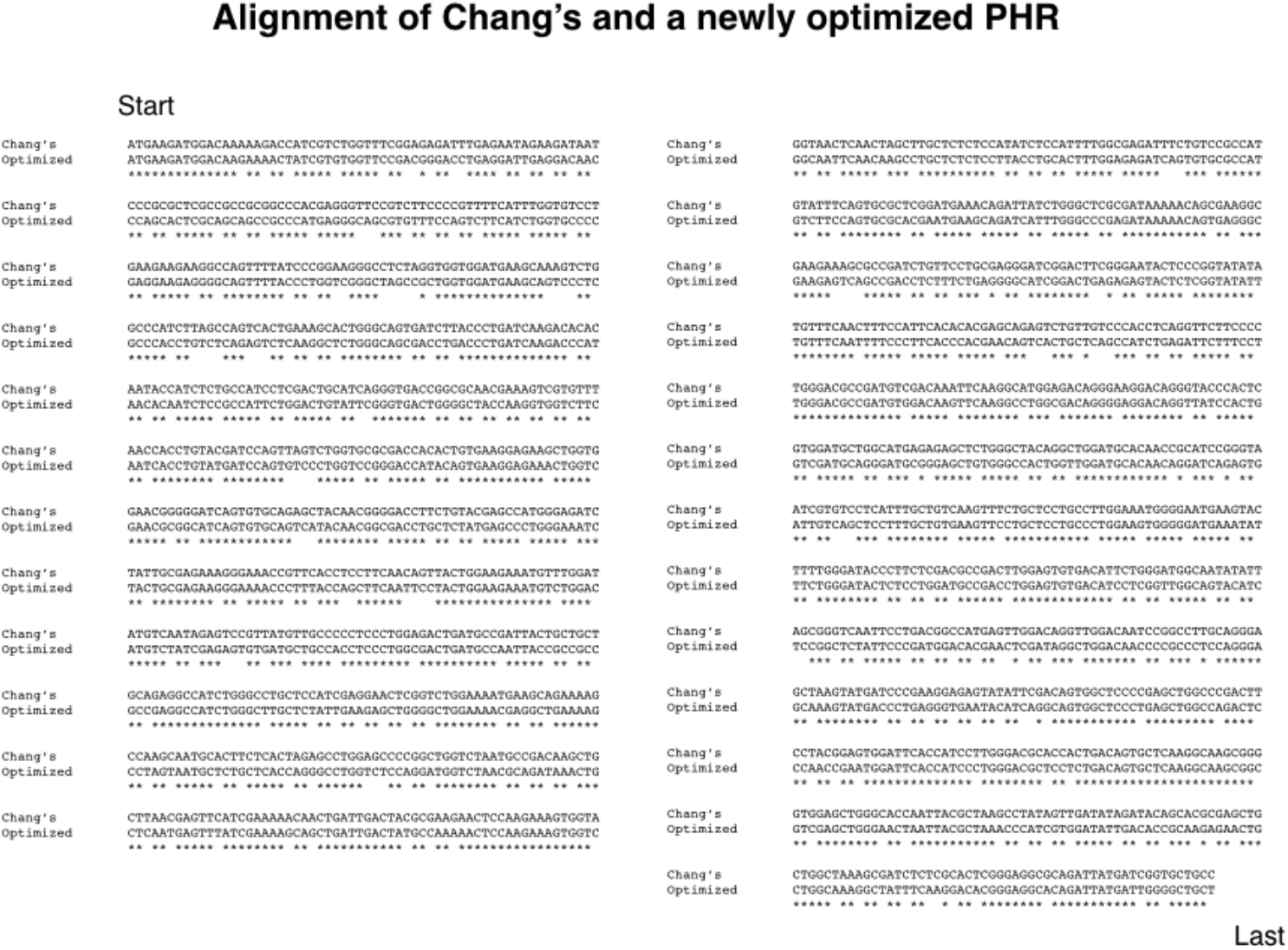
Homology between original PHR and the newly optimized PHR. Homology is 78% at nucleotide level.

**Supplemental Figure 3.**
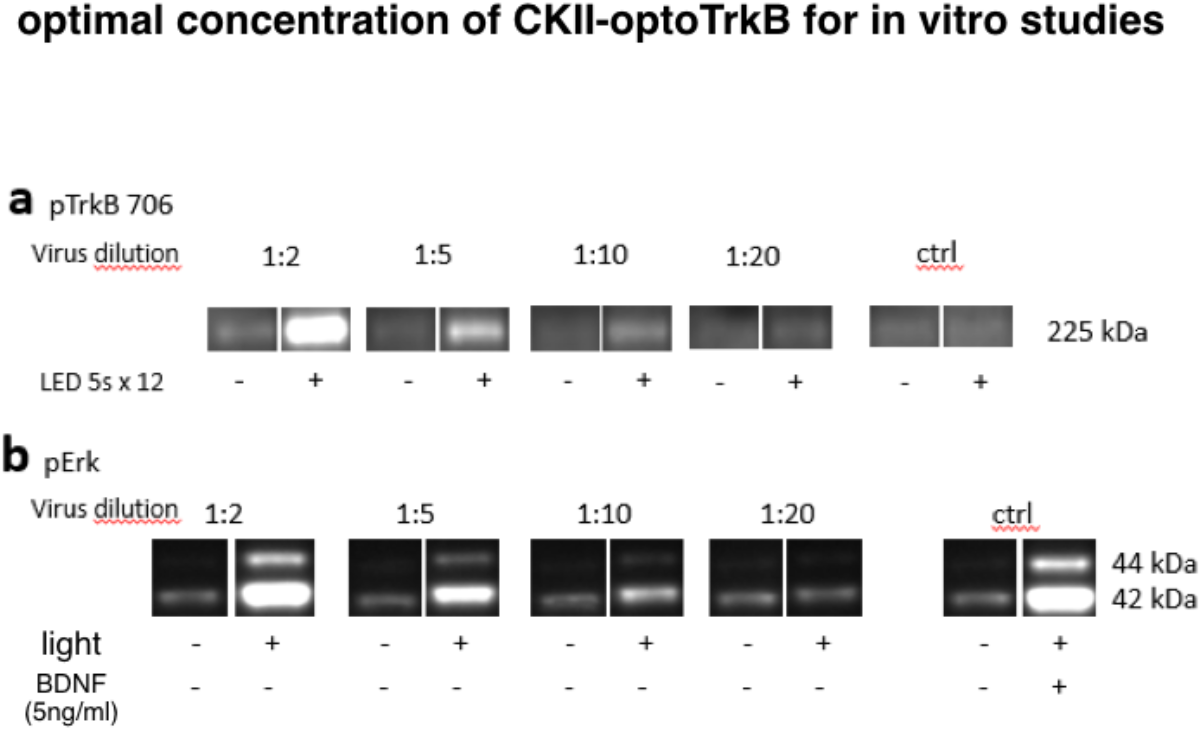
Optimal concentration of CKII-optoTrkB for in vitro studies. Initial virus titer expressed as concentration of p24 protein: 8.37×10^7^pg/ml. (a) Phosphorylation of Y706 site of optoTrkB and (b) phosphorylation of Erk after LED light exposure in primary cortical cells infected with different concentrations of lentivirus. To obtain different concentrations, the virus was diluted in sterile PBS to reach 1:2, 1:5, 1:10 and 1:20 dilutions. The virus was diluted just before administering it to the cells.

**Supplemental Figure 4.**
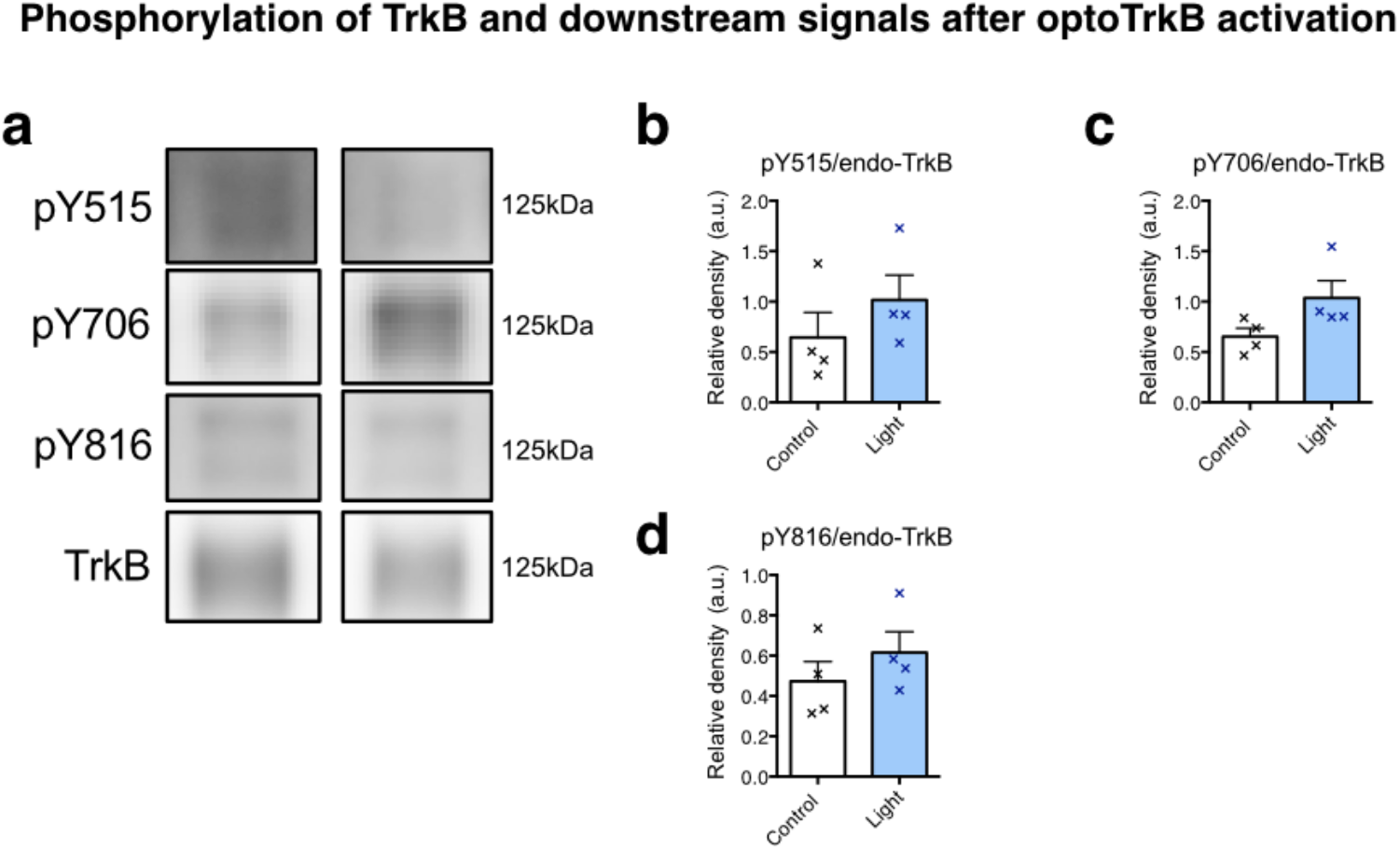
Phosphorylation of endogenous TrkB after light stimulation. (a) Immunoblotting and quantitative analysis of phosphorylation at pY515, pY706, and pY816 sites after light stimulation (12 times for 5 seconds with 1-minute interval) (N = 4, each group). The intensity of endogenous TrkB phosphorylation (125 kDa, left panel) was normalized with the non-phosphorylated version of the same protein. (b-d) There was no significant effect of light exposure on phosphorylation of endogenous TrkB (unpaired t-test: pY515 p = 0.3289; pY706 p = 0.0889; pY816 p = 0.3576).

**Supplemental Figure 5.**
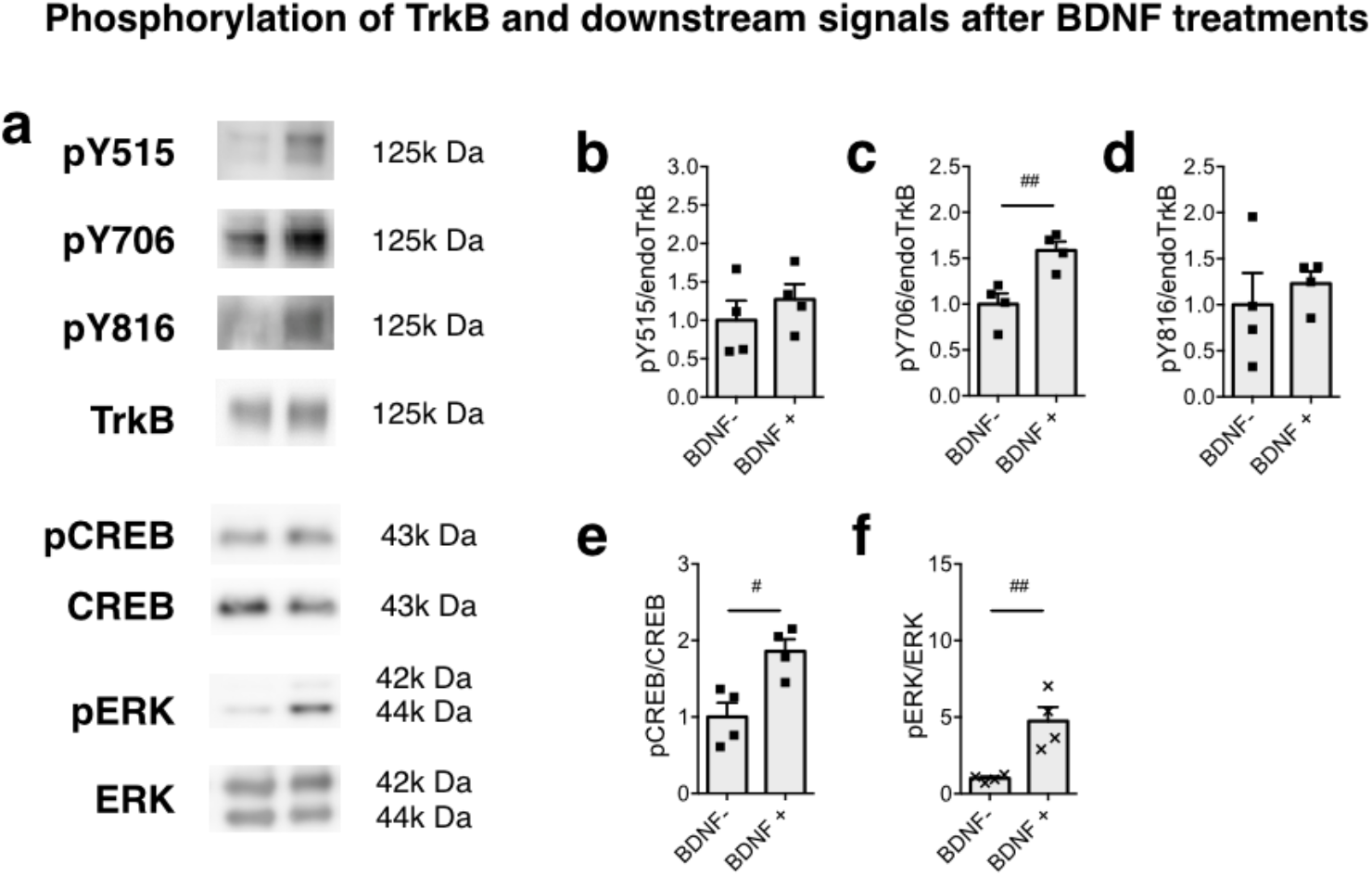
Phosphorylation of TrkB and downstream signals after BDNF treatments in non-infected cultured cortical neurons. (a) Representative images of blotting for phosphorylated and non-phosphorylated endogenous TrkB. the ratio between phosphorylated/non-phosphorylated endogenous TrkB expression at Y515 (b), Y706 (c), Y816 (d), CREB (e), and ERK (f). Unpaired t-test showed that BDNF induced a significant phosphorylation of the Y706 site of TrkB receptor (pY706 p = 0.0084; pY816 = 0.5558; pY515 p = 0.4345) as well as phosphorylation of CREB and Erk (pCREB p = 0.0120; pErk p = 0.0070) # p < 0.05, ## p < 0.01. Bars represent means ± SEM.

**Supplemental Figure 6.**
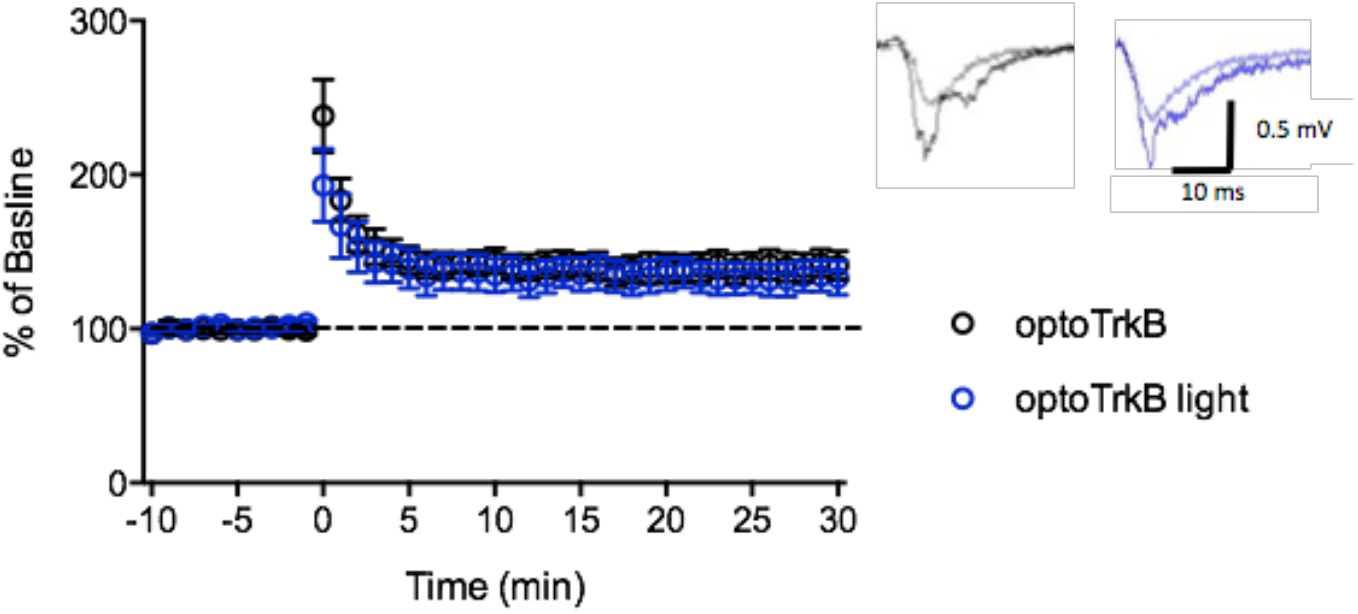
Enhanced neural plasticity after optoTrkB activation ex vivo. The slices from CKII-optoTrkB infected mice were activated by light for 30 seconds. After 30 minutes, a long-term potentiation (LTP) was induced by tetanic stimulation (100 pulses at 50 Hz). At 20 to 30 minutes after tetanization, fEPSPs were larger than baseline in both non-light and light groups and there was no significant difference between the groups (two-way ANOVA, p = 0.0840). Pictures in the right panel show representative traces of fEPSC during baseline and after LTP induction. (optoTrkB, N = 6; optoTrkB light, N = 6). Error bars indicate mean ±SEM.

**Supplemental Figure 7.**
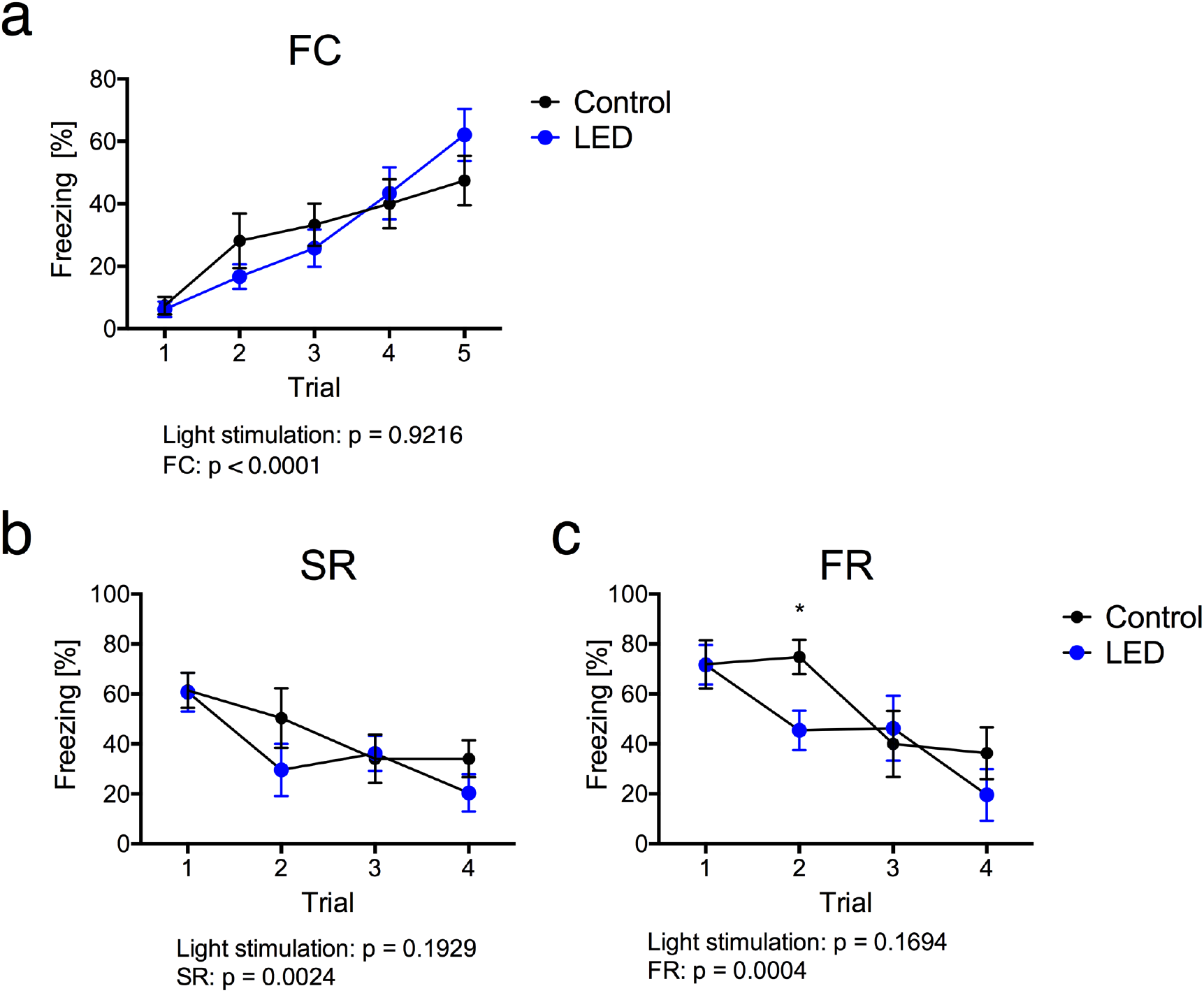
Fear extinction paradigm with mice carrying optoTrkB with extinction training (a) Both control (CKII-optoTrkB infected mice) and light groups (CKII-optoTrkB infected LED-exposed mice) increased freezing during the conditioning/acquisition phase (two-way ANOVA, p < 0.0001) and exhibited the same levels of fear acquisition. Spontaneous recovery (SR) (b) and fear renewal (FR) (c) after activation of CKII-optoTrkB during fear extinction trainings. Previous light stimulation had no effect on freezing duration in SR (two-way ANOVA, p = 0.1929) or in FR (p = 0.1694). There was a significant difference in the 2nd session between control and light-stimulated mice (post hoc: p = 0.0454). N = 8 per group. SR, spontaneous recovery; FR, fear renewal. * p < 0.05, ** p < 0.01. Error bars indicate mean ± SEM.

**Supplemental Fig. 8:**
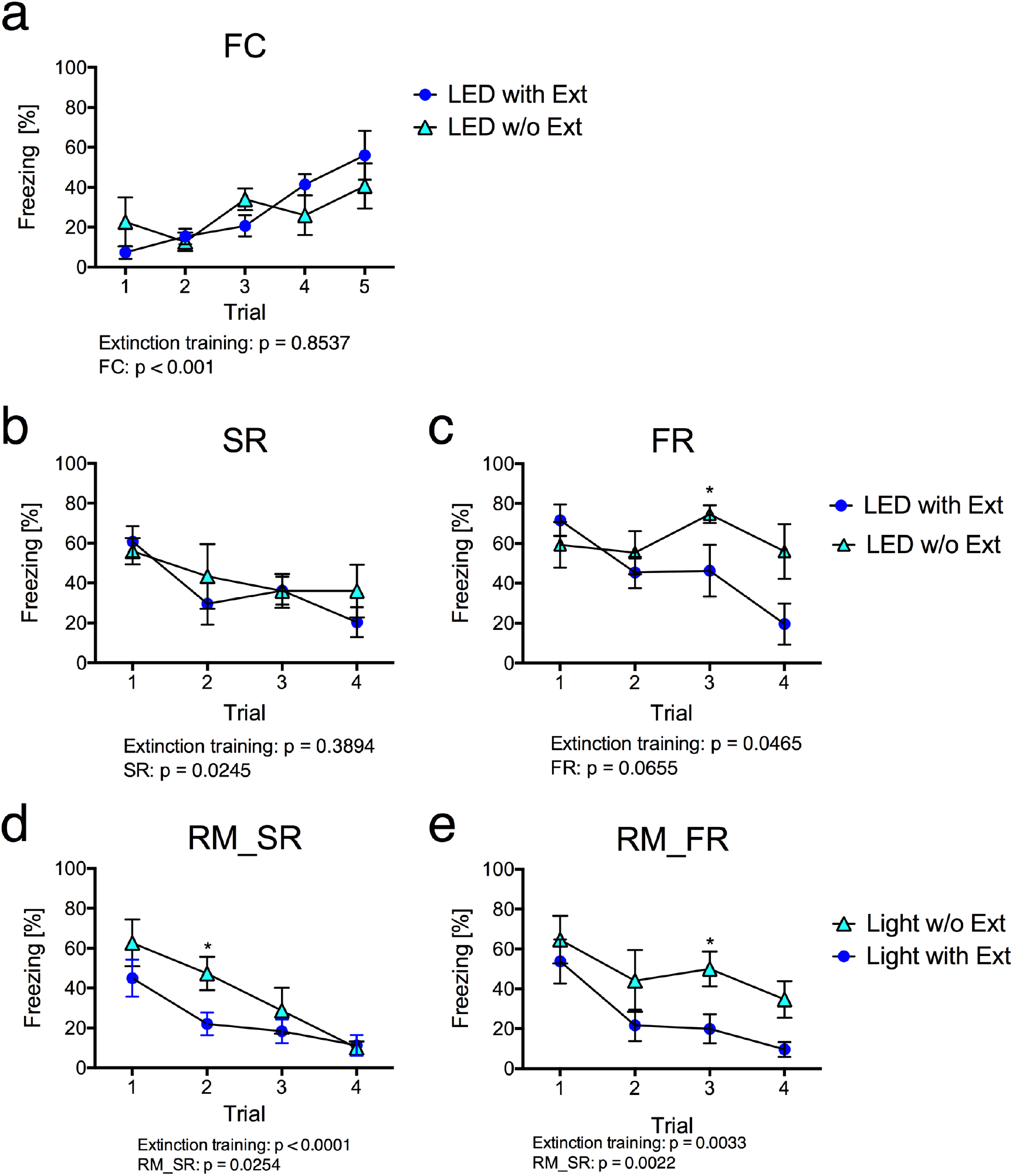
Comparison between groups exposed to light with and without extinction training. (a) Both the light group (CKII-optoTrkB-infected mice stimulated by light with extinction training) and the light group without extinction (CKII-optoTrkB-infected mice stimulated by light without extinction training) increased freezing during the conditioning/acquisition phase (two-way ANOVA, p < 0.001) and exhibited the same levels of fear acquisition. There was a significant decrease in FR (two-way ANOVA, p = 0.0465) (c), but not SR (p = 0.3894) (b) in the group with extinction training compared to the one without extinction training. The light with extinction group showed a stronger decrease in remote spontaneous recovery (RM_SR) (p < 0.0001) (d) and remote fear renewal (RM_FR) (p = 0.033) compared to Light w/o Ext (e). optoTrkB light, N = 8; optoTrkB light w/o extinction, N = 5) * p < 0.05, ** p< 0.01.

**Supplemental table 1.** List of antibodies

| Name | Application | Host | Dilution | Company | Product No |
| --- | --- | --- | --- | --- | --- |
| Horse Radish Peroxidase conjugated Goat Anti-Rabbit | WB | Goat | 1:10000 | BIO-RAD | #1705045 |
| Horse Radish Peroxidase conjugated Goat Anti-Mouse | WB | Goat | 1:10000 | BIO-RAD | #1705047 |
| Mouse anti-GFP | IHC | Mouse | F7 (1:625)<br>C8 (1:667) | Memorial SloanKettering<br>Monoclonal Antibody | 04: clone19F7<br>02: clone19C8 |
| Chicken anti-MAP2 | IHC | Chicken | 1:5000 | Abcam | ab11267 |
| Mouse anti-CAMKII | IHC | Mouse | 1:500 | Abcam | ab22609 |
| Rabbit anti-FosB | IHC | Rabbit | 1:500 | Santa Cruz | sc-7203 |
| Chicken anti-GFP | IHC | Chicken | 1:1000 | Abcam | ab13970 |
| Alexa 546 Goat anti-mouse | IHC | Goat | 1:400 | LifeTechnologies | A21123 |
| Alexa 647 Goat anti-chicken | IHC | Goat | 1:400 | LifeTechnologies | A21449 |
| Alexa 546 Donkey anti-rabbit | IHC | Donkey | 1:500 | Thermo Fisher | A10040 |
| Alexa 647 Donkey anti-mouse | IHC | Donkey | 1:500 | Thermo Fisher | A31571 |
| Alexa 488 Donkey anti-chicken | IHC | Donkey | 1:500 | Jackson | AB_2340375 |

WB: Westernblotting
IHC: Immunohistochemistry

**Supplemental table 2.**
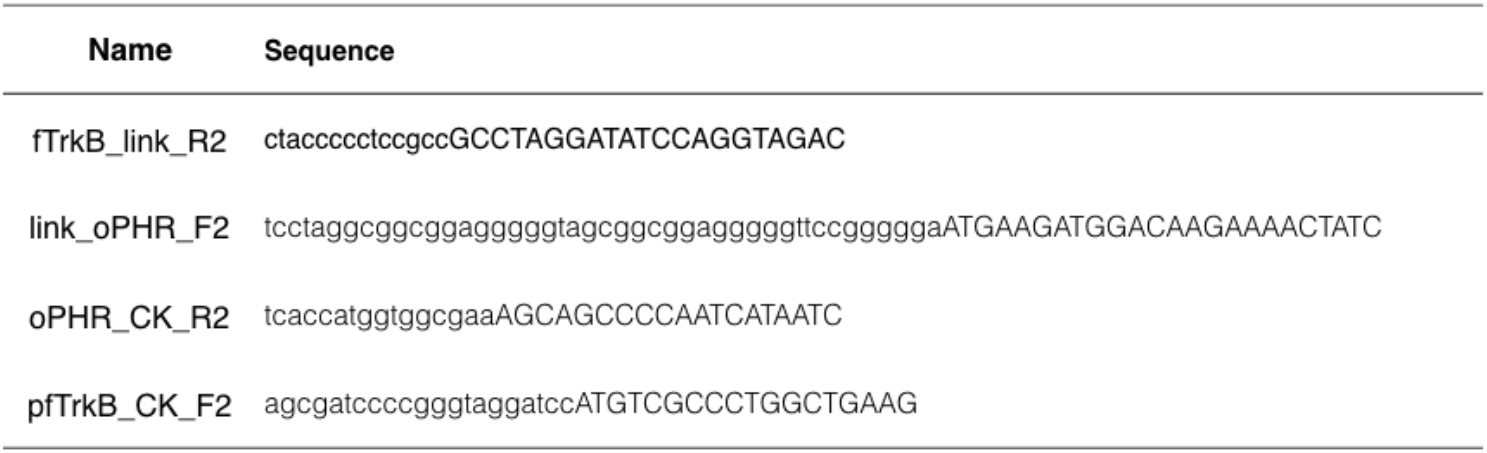
Primers for Gibson cloning

